# *dsSwissKnife*: An R package for federated data analysis

**DOI:** 10.1101/2020.11.17.386813

**Authors:** Iulian Dragan, Thomas Sparsø, Dmitry Kuznetsov, Roderick Slieker, Mark Ibberson

## Abstract

**Summary:** *dsSwissKnife* is an R package that enables several powerful analyses to be performed on federated datasets. The package works alongside DataSHIELD and extends its functionality. We have developed and implemented *dsSwissKnife* in a large IMI project on type 2 diabetes, RHAPSODY, where data from 10 observational cohorts have been harmonised and federated in CDISC SDTM format and made available for biomarker discovery.

**Availability and implementation:** *dsSwissKnife* is freely available online at https://github.com/sib-swiss/dsSwissKnife. The package is distributed under the GNU General Public License version 3.

## Introduction

Many international research projects benefit from the use of pre-existing datasets, which allow increased statistical power for finding relevant and robust associations. This is particularly important for biomedical research collaborations where the goal is frequently to attempt to combine patient data from multiple observational clinical cohorts for better patient stratification and biomarker discovery. There are several problems encountered here. First, there are often legal or ethical constraints that prevent storing individual patient-level data on a central repository for analysis or transferring of data between participating centres. Second, the data are generally not stored in the same format (different variable names and use of non-standard units for measurements), which makes performing the same analysis at different centres problematic. There have been advances in recent years exploring the use of federated database systems, which enables researchers to work on the same version of the data, while the data themselves are kept safe behind local firewalls (Wolfson, et al., 2010).

In order to perform federated analyses, calculi are distributed over a data network and only summary-level data (e.g. partial mean values or regression coefficients) are exchanged, enabling final consolidation of the results. This method has been implemented under the name DataSHIELD (**D**ata **A**ggregation **T**hrough **A**nonymous **S**ummary-statistics from **H**armonised **I**ndividual-lev**EL D**atabases) (Gaye, et al., 2014; Wilson, et al., 2017), which has been recognized by the scientific community and had been used by several groups working in statistical treatment of public health and medical data (Cai, et al., 2017; Doiron, et al., 2013; Doiron, et al., 2017; Pastorino, et al., 2019; Zöller, 2018). It has also been used as an indispensable part for routine data management in several projects in clinical bioinformatics (https://research.ncl.ac.uk/datashield/about/whousesdatashield/).

*dsSwissKnife* is an R package that uses core DataSHIELD functionality for secure communication with remote servers, which also extends its analysis capabilities to include functions for PCA, K-means clustering, cox proportional hazard models, mixed linear models, KNN imputation, Similarity Network Fusion, Adaptive group-regularized ridge regression (GRridge) and Random forest. We have set up a federated database comprising 10 observational clinical cohorts within the IMI project RHAPSODY on type 2 diabetes (T2D) (imi-rhapsody.eu). The RHAPSODY federated database uses CDISC SDTM-formatted Opal (https://www.obiba.org/pages/products/opal/) databases connected to a central server for user administration. *dsSwissKnife* is deployed on this system and is used to perform cross-cohort analyses, whilst the patient-level data remain safely behind local firewalls of their host institutions.

## Results

Ten study populations were harmonized and restructured into a common data model with standardized variable description (CDISC SDTM). Each harmonised dataset was uploaded to a local Opal server hosted by the local institution responsible for the cohort and connected to a central server hosted at Swiss Institute of Bioinformatics (SIB) in Lausanne, Switzerland. The total number of individuals in the database is 47,826 across ten study populations, including various longitudinal anthropometric, clinical as well as disease history and medication related to T2D.

To demonstrate the usability of *dsSwissKnife* for the RHAPSODY federated database, we performed a PCA analysis across all ten study populations with the clinical variables age, body mass index (BMI), high-density lipoprotein (HDL) cholesterol and hemoglobin A1c (HbA1c). *dsSwissKnife* was used to prepare the data from each cohort, scale the clinical variables, perform the PCA analysis and plot the resulting PCA components. Figure 1 shows a biplot of the resulting first and second PCA components stratified into a) study population and b) known to have T2D (Y/N). Although the figure is included here as a proof-of-concept rather than to present analysis results, the scatterplots clearly show differences between cohorts based on these 4 variables. ABOS is an obese diabetic cohort which shows the most separation from the other cohorts in the plot, driven by HbA1c and BMI; GoDARTS, DCS and ANDIS are diabetes progression cohorts and form the main cluster of individuals towards the lower part of the plot; by contrast, the majority of DESIR, CoLAUS, MDC and BOTNIA individuals are non-diabetic or prediabetic, forming a cluster towards the right-hand side of the plot. As expected, the main drivers separating T2D from ND are HbA1c and age (Figure 1B).

**Figure 1.**
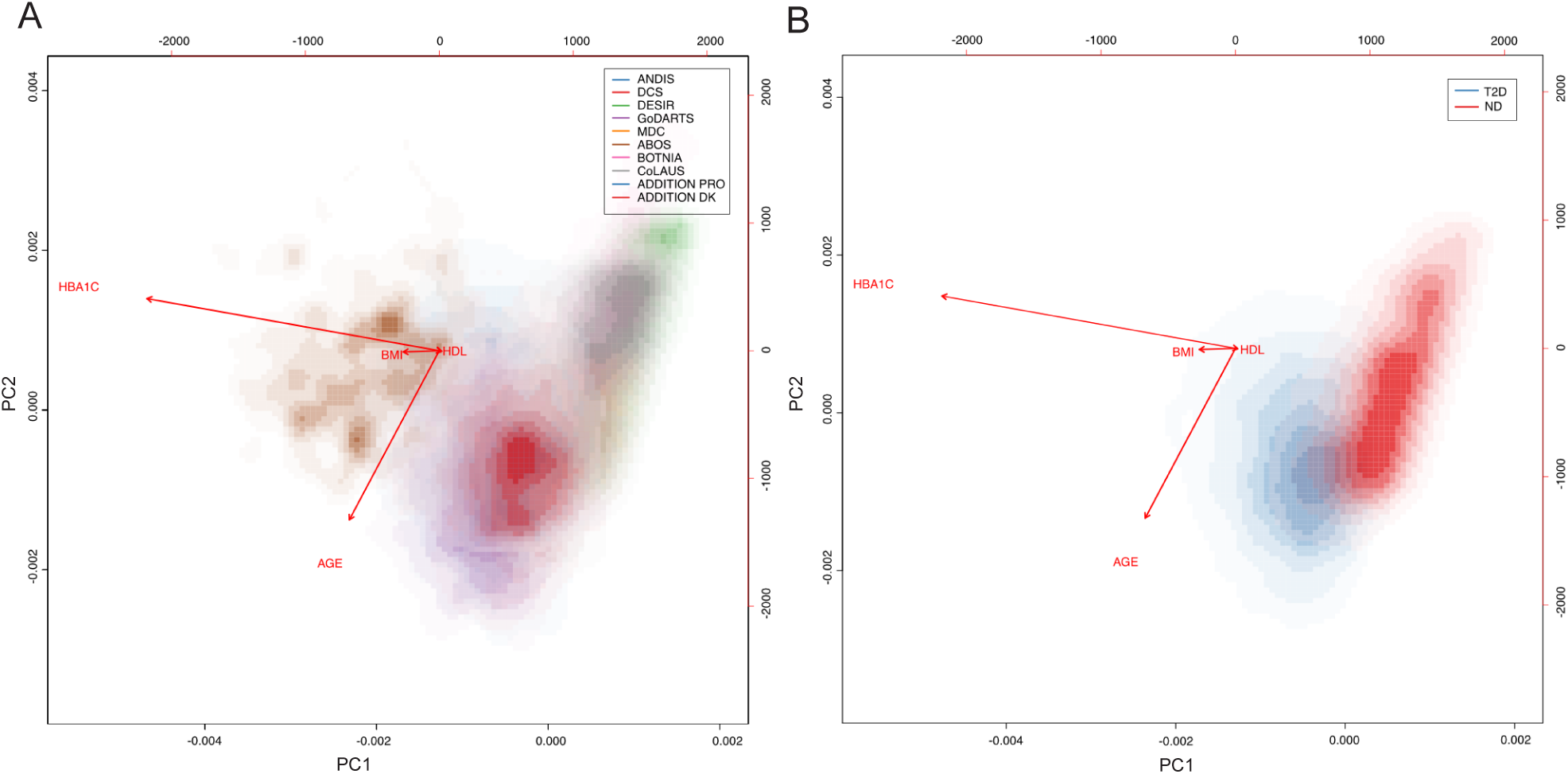
Smooth scatter plots showing the first two principal components in a PCA of a virtual cohort composed of 10 federated cohorts coloured by (A) Federated cohort and (B) T2D status. Stronger colours indicate a higher density of points in an area. Smooth scatter plots are used rather than standard scatterplots to protect patient data. The *dsSwissKnife* function *dssPrincomp* was used for performing the PCA and the function *dssSmooth2d* was used to produce the scatterplots. T2D: type diabetes; ND: non-diabetic.

## Conclusion

The idea of distributed federated analysis is not new; in fact, several initiatives and projects have set up such systems in various research areas, which enable remote analyses to be performed without any copying of individual-level data. Here, we present an R package, *dsSwissKnife*, which extends existing capabilities for federated analysis of sensitive patient data and demonstrate its use in a federated database of 10 clinical observational cohorts. The federated database together with *dsSwissKnife* constitute what is, to our knowledge, the first federated analysis system for the study of diabetes. The setup and development of our system has been driven by the needs and requirements of the RHAPSODY project, with the focus on enabling biomarker discovery. Initially, we were limited by the functionality provided in DataSHIELD, which lacked some of the formatting and statistical methods we required. This gave rise to the development of a new set of tools, which are designed to work alongside DataSHIELD, and also complement and extend it. The resulting *dsSwissKnife* R package can be used with any federated database and includes new computational tools, such as PCA, k-means clustering, KNN imputation and extends the range of linear modelling to include mixed linear models and a faster implementation of linear regression.

Federated analysis similar to the one we present here offers the prospect of providing a scalable solution for access to clinical cohorts in biomarker studies of the future. In such a system, nodes could be shared between different projects and access to each node controlled on a project basis. Any new cohort wishing to participate in such a system would need to provide the computational and data management resources necessary for setting up and maintaining a federated node. In return, they will be able to access detailed clinical data from other federated cohorts in real-time, be able to assess what data are available and to test hypotheses in advance. This would both streamline the discovery process and foster collaboration, helping to maximize the return on investment in future clinical and biomarker research.

## Acknowledgements

Thank you to Nick Giordano, Karina Banasik and Leen ‘t Hart for critical reading of the manuscript and helpful suggestions. This project has received funding from the Innovative Medicines Initiative 2 Joint Undertaking under grant agreement No 115881 (RHAPSODY). This Joint Undertaking receives support from the European Union’s Horizon 2020 research and innovation programme and EFPIA. This work is supported by the Swiss State Secretariat for Education, Research and Innovation (SERI) under contract number 16.0097. The opinions expressed and arguments employed herein do not necessarily reflect the official views of these funding bodies.

## Notes

https://github.com/sib-swiss/dsSwissKnife

### Competing Interest Statement

The authors have declared no competing interest.

